# Passenger mutations can accelerate tumor suppressor gene inactivation in cancer evolution

**DOI:** 10.1101/202531

**Authors:** Dominik Wodarz, Alan C. Newell, Natalia L. Komarova

**Affiliations:** Department of Ecology and Evolutionary Biology, 321 Steinhaus Hall, University of California, Irvine, CA 92697; Department of Mathematics, Rowland Hall, University of California, Irvine, CA 92697; Department of Mathematics, The University of Arizona, 617 N. Santa Rita Ave, Tucson, AZ 85721

## Abstract

Carcinogenesis is an evolutionary process whereby cells accumulate multiple mutations. Besides the “driver mutations” that cause the disease, cells also accumulate a number of other mutations with seemingly no direct role in this evolutionary process. They are called passenger mutations. While it has been argued that passenger mutations render tumors more fragile due to reduced fitness, the role of passenger mutations remains understudied. Using evolutionary computational models, we demonstrate that in the context of tumor suppressor gene inactivation (and hence fitness valley crossing), the presence of passenger mutations can accelerate the rate of evolution by reducing overall population fitness and increasing the relative fitness of intermediate mutants in the fitness valley crossing pathway. Hence, the baseline rate of tumor suppressor gene inactivation might be faster than previously thought. Conceptually, parallels are found in the field of turbulence and pattern formation, where instabilities can be driven by perturbations that are damped (disadvantageous), but provide a richer set of pathways such that a system can achieve some desired goal more readily. This highlights, through a number of novel parallels, the relevance of physical sciences in oncology.

## Introduction

The development and progression of cancer is an evolutionary process whereby cells accumulate multiple mutations, which enables them to break out of homeostasis and to proliferate out of control. The mutations that enable this process typically confer a selective advantage to cells and have been called driver mutations [1–5]. Tumors, however, are highly heterogeneous and cells also contain a variety of other mutations, called passenger mutations [1–5]. They arise from random mutations in sequences that do not contribute directly to disease, facilitated by exposure to mutagenic processes and lack of repair [6]. While passenger mutations have been thought to have minimal biological consequences on the disease process, the properties and role of passenger mutations remain poorly understood. Recent data indicate that passenger mutations carry a certain fitness cost [7,8], and that they might therefore render tumors more fragile, which could be exploited therapeutically [4,7,9]. One particular evolutionary process that is central to carcinogenesis and cancer progression in the inactivation of tumor suppressor genes (TSG) ^9,10^. This typically requires two mutational hits because both copies of the gene need to be inactivated to achieve full loss of function [10]. Different tumor suppressor genes display different characteristics, and in principle, the inactivation of only one copy of the gene either results in no change in the fitness of the cell, or it could entail a certain selective disadvantage. To inform model assumptions, we will specifically consider the tumor suppressor gene APC, which becomes inactivated early in the development of colorectal cancer [11]. In this case, data indicate that heterozygous APC+/− cells can experience reduced fitness, which means that a fitness valley has to be crossed for the inactivation of APC to occur. Experiments with colorectal cancer cell lines revealed that a truncating mutation in APC has a dominant effect resulting in a spindle checkpoint defect, aneuploidy, and a reduced proliferation rate of cells [12,13]. Similar effects have been found in vivo in APC^Min/+^ mice [14], which have an APC+/− germ line mutation. In general, if a copy of a tumor suppressor gene is lost as a consequence of aneuploidy, the cell is likely to suffer a fitness reduction (see e.g. references [15,16]). Motivated by these studies, our paper investigates the effect of passenger mutations on the evolutionary dynamics of tumor suppressor gene inactivation, assuming that a fitness valley needs to be crossed.

## Results

We consider a computational model for the inactivation of tumor suppressor genes (TSG) [17,18], where cells only acquire an advantage once they have accumulated two separate mutations, but are neutral or disadvantageous in the presence of only one of the mutations. For convenience, cells with both copies of the TSG present are referred to as TSG^+/+^, and cells with one or both copies of the TSG inactivated are referred to as TSG^+/−^ and TSG^−/−^ respectively. Much evolutionary work has been performed that studied how fast such fitness valleys/plateaus can be crossed, depending on the scenario under consideration [19–30]. To study the role of passenger mutations, we employ a stochastic agent-based model that is also referred to as a contact process. This model assumes the existence of N spots, which can either be empty or contain a cell. Each time step, the system is sampled M times, where M is the number of cells present. If the chosen spot contains a cell, it can divide and die with defined probabilities. When a cell is chosen to divide, a target spot is chosen randomly from the whole system, and division only proceeds if this target spot is empty. Upon division, mutations can occur that give rise to different cell genotypes (Figure 1A). TSG+/+ cells without passenger mutations are denoted by *x* and attempt division with a probability L_x_ per cell per update. TSG+/− cells without passenger mutations are denoted by *y* and have a fitness cost of s_1_ (s_1_ ≤1), such that their division probability is s_1_ L_x_. TSG+/+ cells that also contain passenger mutations are denoted by z and have a fitness cost s_2_ (s_2_<1). TSG+/− cells that also contain passenger mutations are denoted by w and have a fitness cost s_3_ (s_3_<1). Both TSG+/− populations, y and w, can give rise to the advantageous TSG−/− double mutant. All the mutation processes are defined in Figure 1A]. For simplicity, each cell type is assumed to die with the same rate D. The model was simulated repeatedly, and the fraction of realizations when an advantageous TSG^−/−^ mutant had been generated by a defined time threshold was determined. We compared simulations without passenger mutations (n=0) with those that did allow the generation of passenger mutations (n>0). Two different regimes have been observed [24]: In one regime, the double-hit mutant arises without the intermediate TSG+/− mutant reaching fixation, a process called stochastic tunneling. The second regime can be called sequential fixation, were the intermediate TSG+/− mutant fixates before the double mutant is created.

**Figure 1:**
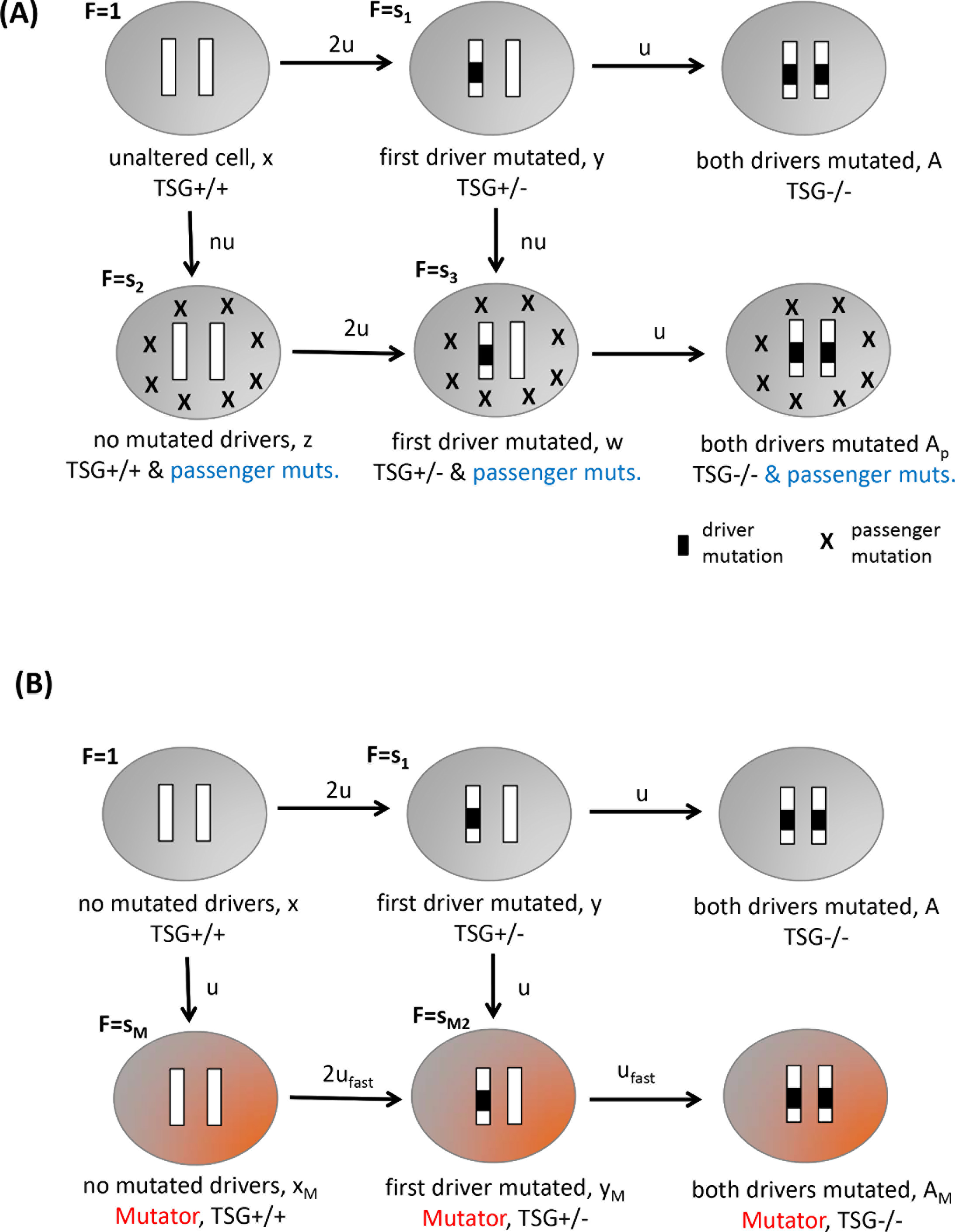
Schematic representation of computational models. (A) Model of TSG inactivation in the context of passenger mutations. The cell types and associated fitness values (F) are explained in the main text. The arrows show the mutational steps that generate the different cell types. The top row of cell types depicts standard evolutionary processes where the two copies of the TSG are sequentially inactivated. In addition, the model assumes that with a rate nu, cells can accumulate passenger mutations. Cells with passenger mutants can also inactivate the TSG, as shown. (B) Model of TSG (APC) inactivation in the context of Lynch Syndrome and mutator phenotypes in colorectal cancer. This model does not contain passenger mutations. The basic evolutionary processes (along the top row of cells) are the same as above. The difference is that unmutated TSG+/+ cells can inactivate mismatch repair mechanisms with a rate u, giving rise to mutator phenotypes that are characterized by an elevated mutation rate u_fast_.

For the tunneling regime, we find that the presence of passenger mutations can accelerate the generation of the advantageous double mutant (Figure 2Ai). The magnitude of this effect increases with higher fitness of the passenger mutants, s_2_ (Figure 2Ai). This requires that (i) the number of passenger mutations that can be accumulated, n, is sufficiently large relative to the inverse of the mutation rate and (ii) the intermediate TSG^+/−^ mutation reduces the fitness to a lesser degree in cells with passenger mutations than in cells without passenger mutations, i.e. there are epistatic interactions between drivers and passengers [31,32]. The exact condition is *S*_*3*_>*S*_*1*_*S*_*2*_, see Supplementary materials for computational details. This is a necessary condition for the passenger mutations to accelerate evolution (both in the tunneling regime and in the sequential fixation regime, see below). In the Discussion section, we describe a specific example where the fitness of TSG+/− mutants is context dependent, indicating that an assumed occurrence of epistasis in such cells is biologically relevant.

**Figure 2:**
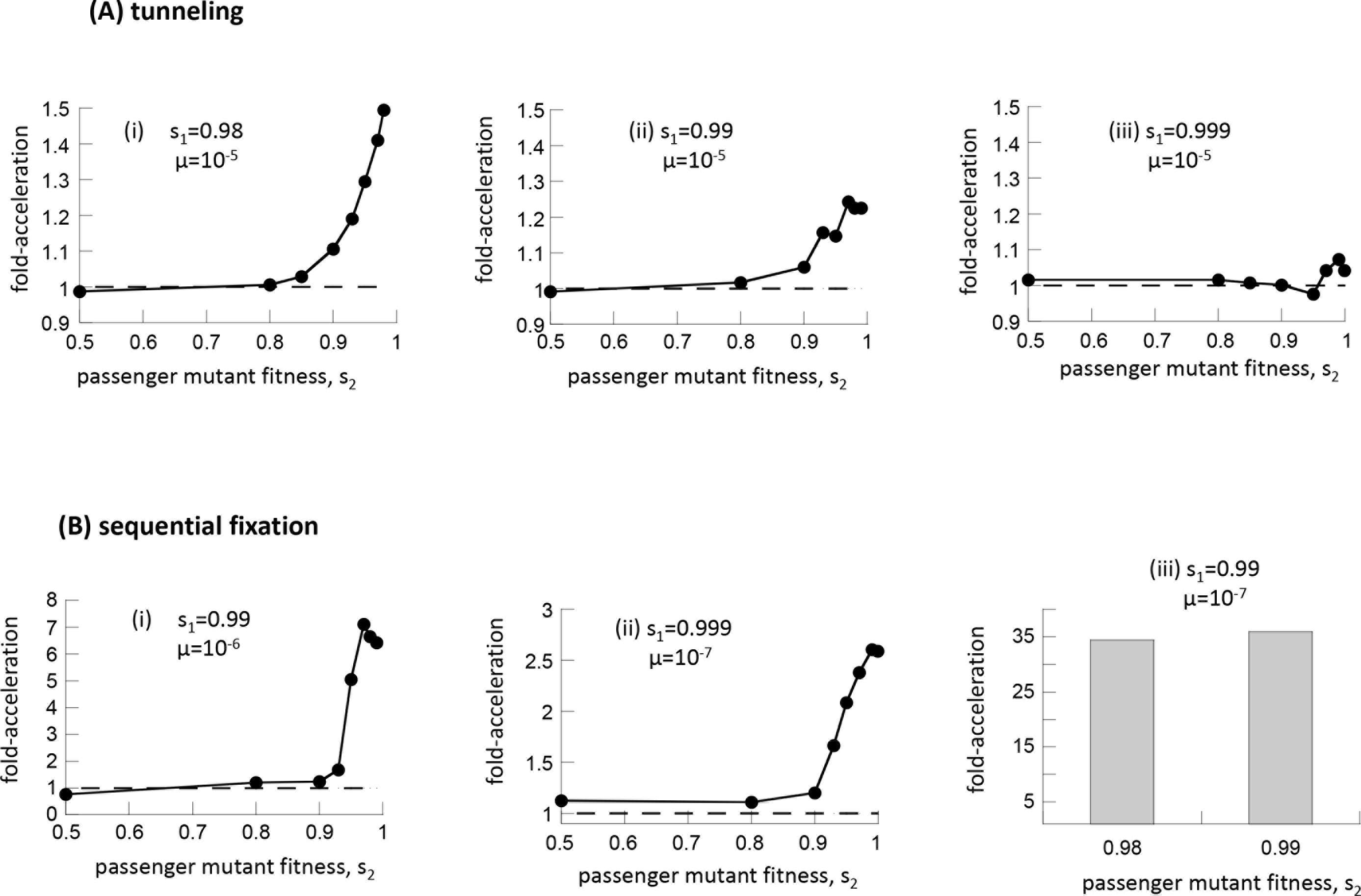
Accelerated crossing of fitness valleys (i.e. inactivation of TSGs) in the presence of passenger mutations. The computer simulations were run repeatedly, and the fraction of realizations in which the double mutant was created before a time threshold T_thr_ was determined. The number of simulations / sample sizes required by rare events is large and were chosen using reference [45]. For a fixed margin of error, the more rare the events, the larger the sample size. The margins of errors of our study are acceptable, as even in the worst situation (N=543449, recorded fraction=0.000077), the margin of error is less than one third of the estimated proportion. Each graph plots the “fold acceleration”, which is the fraction of runs where the double mutant was generated in the presence of passenger mutations with fitness s2, divided by the same measure in the absence of passenger mutations. The Z-score for population proportions was used to determine whether the difference in outcome between simulations with and without passenger mutations was statistically significant (the distribution under the null hypothesis, when the two true proportions are the same, is asymptotically normal). (A) Parameter regime where the double mutant evolved through a tunneling pathway. Panels (i-iii) show that a reduced cost of the intermediate TSG+/− mutant (higher value of s_1_) leads to a reduced effect of passenger mutations on the rate of evolution. Differences between outcomes with and without passenger mutations were statistically significant for (i) s_2_≥0.93, (ii) s_2_≥0.93, and (iii) s_2_=0.99. (B) Parameter regime where the double mutant evolves by sequential fixation, in which passenger mutants have a stronger accelerating effect on the rate of evolution. Differences between outcomes with and without passenger mutations were statistically significant (p<0.05) for (i) s_2_≥0.95, (ii) s_2_≥0.93, and (iii) s_2_=0.98 & s_2_=0.99. In panel (iiii), the fold acceleration was only determined for the highest values of s_2_ (where the effect is strongest), due to the extensive computational cost associated with this parameter set. Remaining parameters were chosen as follows. *L*_*X*_=*0.15*, *D*=*0.01*, *s*_*3*_=*s*_*2*_. *n*=*5×10^−2^/μ*, *N*=*2500*. The results do not depend on the assumption s_3_=s_2_. How s_3_ needs to depend on s_2_ and s_1_ for the results to hold is defined in the supplementary materials. The time thresholds are given as follows for the individual graphs. (A) (i) *T*_*thr*_=*8,000*; (ii) *T*_*thr*_=*8,000*, (iii) *T_thr_*=*5,000*; (B) (i) *T*_*thr*_=*120,000*; (ii) *T*_*thr*_=*1,200,000*; (iii) *T*_*thr*_=*1,500,000*.

The reason for the accelerated evolution in the presence of passenger mutations is that these mutations increase the relative fitness of the intermediate TSG+/− cells through a set of complex interactions. If the wild-type population consists mostly of cells without passenger mutations, the evolutionary dynamics are largely driven by the x, y system, which is relatively slow due to the more pronounced disadvantage of y. The larger the proportion of cells with passenger mutations, however, the more the evolutionary dynamics are driven by the z, w system, where the intermediate mutant suffers an overall lower fitness cost. This allows the total population of intermediate TSG^+/−^ cells to persist at a higher selection-mutation balance, making it more likely to generate the double mutant. The average rate of double mutant generation can be calculated from ordinary differential equations (ODEs, see Supplementary Materials), which is a reasonable model that quantifies population dynamics in the tunneling regime. The rate of double mutant generation is increased by the presence of passenger mutants, with more pronounced effects for larger values of s_2_ (Figure 3), thus explaining our observations. If the fitness of passenger mutants crosses a threshold (which depends on the total rate of passenger mutant generation), the cells with passenger mutations, z, outcompete those without passenger mutations, x (because z is generated by x). In this regime, the advantageous double mutants are created fastest because the intermediate TSG+/− mutants (w) have the highest relative fitness out of all scenarios. This might be a biologically relevant parameter region given the ubiquitous occurrence of passenger mutations in cancer cells, and even in aged non-cancerous tissue [33,34]. We note that in this model, the generation of cells with passenger mutations does not increase the total population size and hence does not provide additional targets for mutation. The accelerating effect of passengers stems from the overall reduction in population fitness and the consequent elevation of the relative fitness of intermediate TSG+/− mutants.

**Figure 3:**
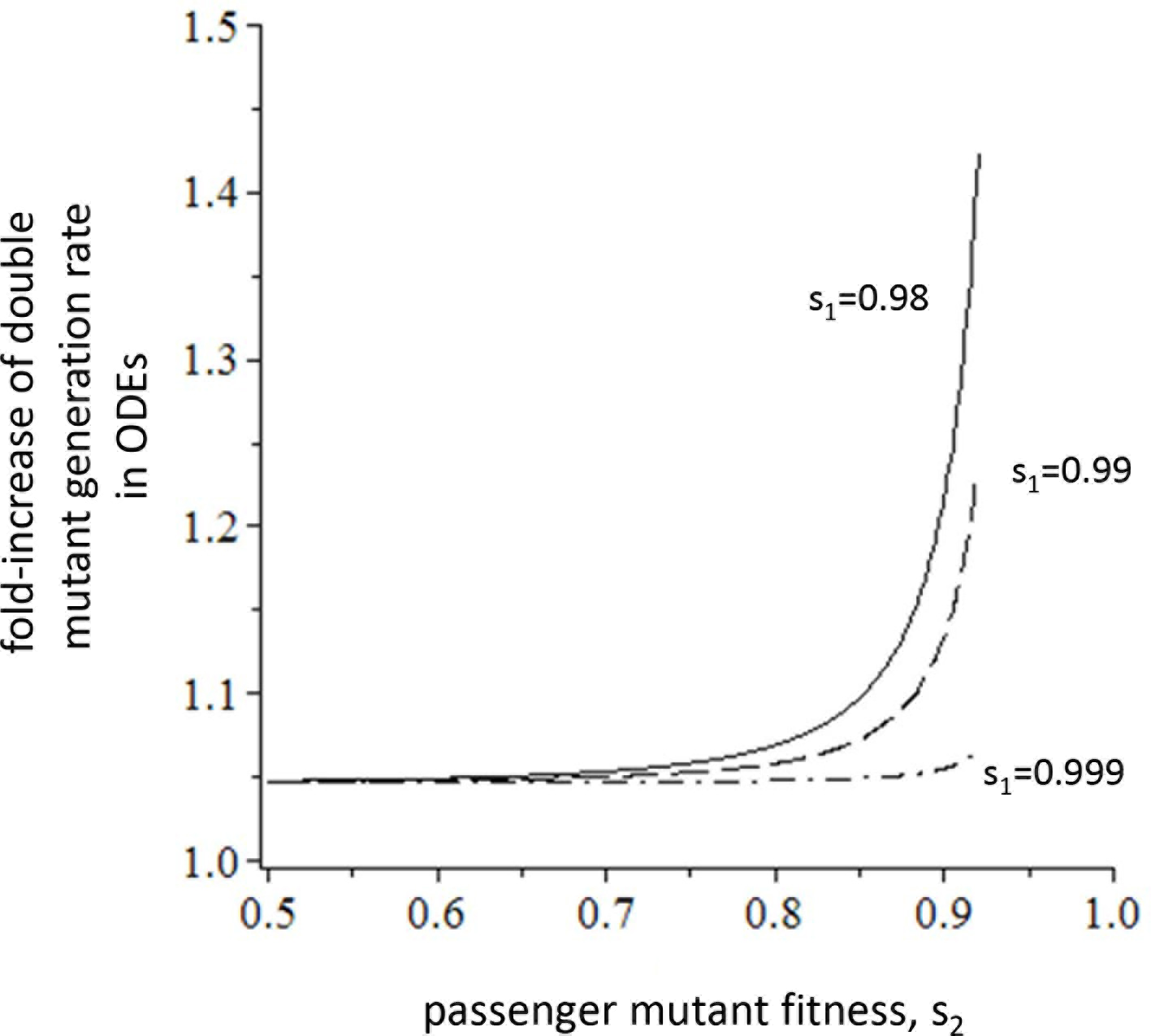
The average behavior of the contact process can be described by ordinary differential equations given in the Supplementary Materials. From these ODEs, equilibrium populations sizes, and hence the average rate of double mutant generation at equilibrium, can be calculated. In the presence of passenger mutations, this is given by (s_1_y*+ s_3_w*)[1−(x*+y*+z*+w*)/k], where * denotes equilibrium population sizes. This is divided by the rate of double mutant generation in the absence of passenger mutations, given by s_1_y*[1−(x*+y*)/k]. This yields the fold increase of the double mutant generation rate that is mediated by passenger mutations, and is plotted in the graph for different values of s_1_ (fitness cost of TSG+/− cells without passenger mutations, y). Parameter values were chosen as follows: *r*=*0.1*, *d*=*0.01*, *μ*=*10*^−5^, *n*_*1*_=*2*, *n*_*2*_= *5000*, *k*=*2500*.

The extent to which passenger mutants accelerate fitness valley crossing further depends on the relative fitness of intermediate TSG^+/−^ mutants without passenger mutations (s_1_, the fitness cost of population y). The closer the fitness of the intermediate mutant y-population is to the fitness of the wild-type x-population (s_1_⟶1), the less pronounced the accelerating effect (compare Figures 2Ai-iii). This is also seen in the ODE predictions, which show that for larger values of s_1_, the average rate of double mutant generation is accelerated by passenger mutations to a lesser extent (Figure 3). The reason is that for higher s_1_, the intermediate TSG^+/−^ mutants (y) have less of a disadvantage compared to population x, which leaves less room for improvement by the z, w interactions. Thus, if the intermediate TSG+/− mutant is almost neutral with respect to the wild-type, passenger mutants are not likely to accelerate evolution in the tunneling regime.

Next, consider the parameter regime where the intermediate mutant fixates prior to the generation of the double mutant (sequential fixation). This tends to occur for parameters where the generation of the double mutant takes a longer period of time due to lower mutation rates or smaller population sizes. In this scenario, the accelerating effect of passenger mutations can be significantly more pronounced than in the tunneling regime (Figure 2B). For a physiologically realistic rate of gene inactivation (10^−7^ per gene per division), even if the TSG+/− mutants without passenger mutations only have a 0.1% fitness cost (s_1_=0.999), and if the passenger mutations lead to a 1% fitness cost (s_2_=0.99), the presence of passenger mutations can accelerate the emergence of the double mutant almost 3-fold (Figure 2Bii). If the fitness cost of the intermediate TSG+/− mutant is 1%, the acceleration can be up to 35-fold (Figure 2Biii). The reason is that the fixation probability of the TSG+/− mutants is markedly higher when the dynamics are governed more by the z,w system compared to the x,y system.

These results remain robust if instead of assuming that all passenger mutants have the same fitness cost, those fitness cost values are taken from a power function distribution between zero and one, with averages given by s_2_ and s_3_. This potentially allows for some significantly deleterious passenger mutants even though many of them can be close to neutral (Supplementary Materials). Results are further shown to remain robust in a spatially explicit model, where dividing cells place their offspring in a randomly chosen spot nearby (Supplementary Materials). Finally, the same patterns are observed in a constant population Moran process, which represents tissues where normal cells are maintained at carrying capacity and their homeostatic turnover is driven by cell death (Supplementary Materials).

While our models have shown that the presence of passenger mutations can accelerate the rate of TSG inactivation, the question arises how significant this acceleration can be. To gauge that, we compare the degree of acceleration that can be observed in our passenger mutation model to the acceleration observed in the context of a different and unrelated process that occurs in colorectal carcinogenesis, and that is known to lead to clinically significant accelerations in evolutionary processes: tumor initiation in Lynch Syndrome patients. It is known that Lynch Syndrome patients develop colorectal tumors with a significantly faster rate than the general population. This is because Lynch Syndrome patients are characterized by a germ line mutation in one copy of a mismatch repair (MMR) gene. Hence a single point mutation can frequently generate MMR-deficient cells that promote mutant accumulation and hence the inactivation of APC. Therefore, we describe tumor formation in the context of Lynch Syndrome (without passenger mutations) in the same kind of computational framework studied so far (Figure 1B), and determine the extent to which evolution is accelerated in that model. If the degree of acceleration observed in the Lynch Syndrome model is of a similar magnitude as the acceleration observed in the passenger model, then there is indication that passenger-induced acceleration can be clinically highly relevant. If the acceleration in the passenger model is much less than that in the Lynch Syndrome model, then the passenger-induced acceleration is less relevant. In particular, we investigated by how much the mutation rate in the mismatch repair (MMR)-deficient cells has to be increased to obtain a degree of evolution acceleration that is comparable to that observed with passenger mutations. To do so, we assumed that genes are inactivated with a rate of 10^-7^ per division, and that an intermediate TSG+/− cell carries a 1% fitness cost, consistent with data that documented reduced growth of APC+/− cells [12]. As before, passenger mutants were also assumed to carry a 1% fitness cost. If we assume that MMR-deficient cells do not carry a fitness cost, we obtain that MMR-deficient cells need to have a 100-500 fold increase in their mutation rate to accelerate evolution to a similar degree as seen in corresponding passenger mutant simulations (Figure 4a). If MMR-deficient cells have a 1% fitness cost, then the fold increase in the mutation rate has to be 1000-5000 fold to match the acceleration afforded by the presence of passenger mutations (Figure 4b). Because this increase in mutation rate is thought to be typical for mismatch-repair deficient colorectal cells [35], this suggests that passenger mutations can have an accelerating effect that is similar in magnitude to acceleration in Lynch Syndrome, pointing to potentially strong biological relevance. If APC+/− cells are characterized by a significantly lower fitness cost, or if the assumed epistatic interactions between drivers and passengers are significantly weaker, this effect would be reduced.

**Figure 4:**
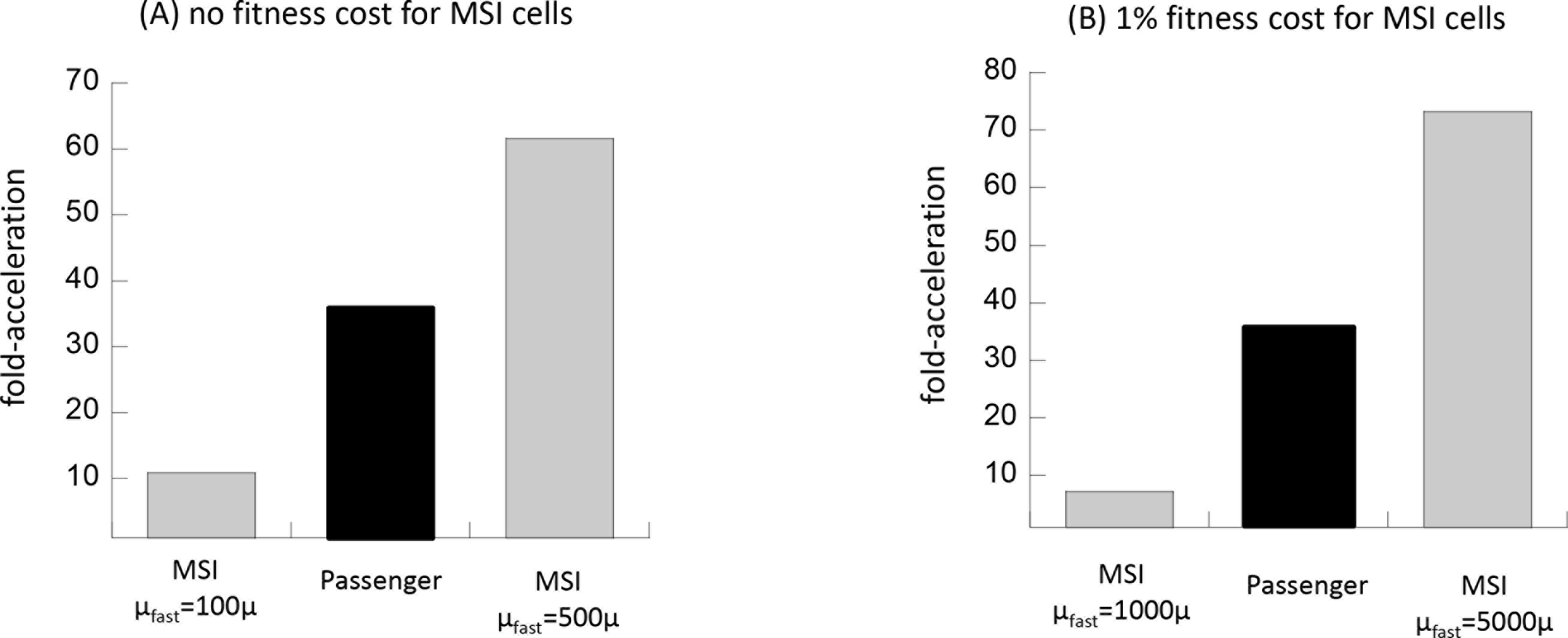
Rate of TSG inactivation in two types of simulations: assuming Lynch Syndrome, which involves the acquisition of microsatellite instability or MSI, i.e. cells with a faster mutation rate (Figure 1B); and assuming the passenger mutant pathway (Figure 1A). The computer simulations were run repeatedly, and the fraction of realizations in which the double mutant was created before a time threshold T_thr_ was determined. For the Lynch Syndrome model, simulations with different accelerated mutation rates, u_fast_, were run. The fraction of runs that resulted in double mutant generation were divided by the fraction obtained without the existence of mutator phenotypes (MSI cells), which is the fold-acceleration depicted by the gray bars. The black bar shows the fold-acceleration derived from the passenger mutation model (without mutator cells), but with otherwise identical parameters. This was done in two settings (A) assuming that mutator cells do not suffer from a fitness cost; (B) assuming that mutator cells are characterized by a 1% fitness cost, brought about by the frequent generation of deleterious mutations. The parameters were chosen as follows. *L*_*x*_=*0.15*, *D*=*0.01*, *s*_*1*_=*0.99*, *S*_*2*_=*0.99*, *S*_*3*_=*0.99 μ*=*10*^*−7*^. For (a) *s*_*M*_=*1*, *s*_*M2*_=*1*. For (b) *S*_*m*_=*0.99*, *S*_*M2*_=*0.99*, *n*=*5×10*^*−2*^/*μ*, *T*_*thr*_=*1,500,000*, *N*=*2500*.

## Discussion and Conclusion

Previous work reported that the presence of passenger mutations can make tumors more fragile in certain circumstances due to a reduction in overall fitness [4,7,9]. Here we have shown that in the context of fitness valley crossing, costly passenger mutations can actually accelerate evolution because they reduce overall population fitness and thereby provide an environment in which intermediate TSG+/− mutants enjoy a higher relative fitness. Although passenger mutations are selected against, their accumulation (even at low numbers) provides access to additional pathways to cancer where the fitness valley is shallower and easier to cross. It has been previously suggested that the process of carcinogenesis could be promoted through a reduction of overall population fitness due to aging and other insults, thus providing a more favorable fitness landscape for the evolution of malignant cells [36]. Our passenger mutations model fits well into this concept.

The models further indicate that this can result in an acceleration of evolution that can be comparable to that observed in patients with a predisposition to genetic instability. This suggests that the “baseline” rate of TSG inactivation in the absence of genetic predisposition and genetic instability can be significantly faster than previously thought, which might be conceptually important for understanding the ability of cells to accumulate a number of carcinogenic mutations in a relatively short period of time [37]. This applies not only to tumor progression, but also to cancer initiation in healthy tissue, which has been shown to contain a significant number of passenger (and driver) mutations [34], especially at advanced age [33].

Our analysis identified possible epistatic interactions between driver and passenger mutations [31,32] to be important for the reported dynamics, and the literature supports this notion. Indeed, it has been pointed out that the classification of mutations into passengers and drivers might be an over-simplification, because the fitness of a given cancer phenotype can be context-dependent [38]. More specifically, we turn again to the tumor suppressor gene APC in colorectal carcinogenesis. There are mouse strains that are heterozygous for the APC^Min^ (multiple intestinal neoplasia) mutation, called APC^Min/+^ mice. They frequently develop intestinal tumors [39,40]. Significant variation in tumor incidence occurs among APC^Min/+^ mice with identical APC mutations and which are kept under identical laboratory conditions [39,40]. This variation is caused by differences in the genetic background of the APC mutation, which in turn depends on variation in “modifier genes” in different mouse strains [39,40] [41]. These are not involved directly in the process of carcinogenesis, but modify the phenotypic properties of APC+/− cells. This indicates that passengers can modulate the fitness of TSG+/− cells, as required by our model to observe accelerated evolution.

Our work adds to the growing literature that investigates the dynamics of fitness valley crossing under various conditions [19–21,23,25–28,42–44]. Beyond this immediate discipline, however, it is also interesting to consider our results in a wider scientific sense. In the presence of passenger mutations, cellular evolvability is predicted to increase through the introduction of disadvantageous cells. The presence of these disadvantageous cells lowers overall population fitness, allowing intermediate TSG+/− mutants to have an overall higher relative fitness, which promotes faster generation of the TSG−/− double mutant. Studies of the onset of turbulence as well as pattern forming systems have revealed mechanisms of instability that act in a very similar manner. For example, in turbulence it has been shown that three-dimensional perturbations on parallel shear flows are damped more strongly than two-dimensional ones; but because of the slow decay, they provide a new base flow on which a new and richer class of fluctuations can grow more rapidly (details in Supplementary Materials). This provides a fundamental connection between principles in the physical sciences and the particular oncology question under consideration.

## Acknowledgement

This study was funded in part by NIH grant U01CA187956.

## 1 ODE description

### 1.1 Logistic model

We used the contact process as a basic model to study the effect of passenger mutations on the rate of fitness valley crossing, i.e. on the emergence of a *TSG*^−/−^ mutant. The contact process is a stochastic agent-based model and is described in the main text. Because the cell populations are assumed to be well-mixed (mass action), the average behavior of the contact process can also be studied by means of the ordinary differential equations. Denote by *x*, *y*, *z*, and *w* the wild type cells, TSG^+/−^ cells, TSG^+/+^ cells with passenger mutations, and TSG^+/−^ cells with passenger mutations respectively. The replication rate of wild type cells is denoted by *r* and the death rate by *d*. The fitness values for these types of cells are given by 1, *s*_1_, *s*_2_, and *s*_3_ respectively. Let us denote by *W* the term describing the competition of the cell types:

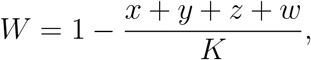

where *K* has the meaning of carrying capacity. Then the ODEs are given by

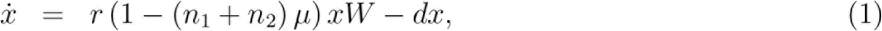

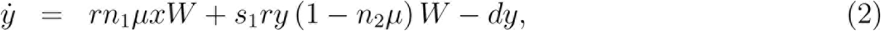

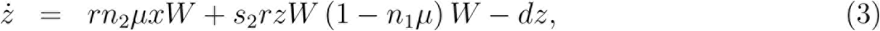

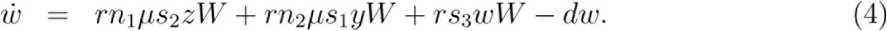

By rescaling time we can assume that *r* = 1 for the purpose of the analysis below (the value of *d* is rescaled accordingly). The replicative cost of the various mutants are given by *s*_1_, *s*_2_, and *s*_3_, which can have values between zero and one. We further assume that

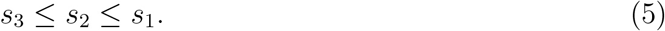

The death rate of cells is given by *d* and for simplicity is assumed to be identical for all cell populations. The number of different passenger mutations that a cell can accumulate is given by *n_1_*, and the number of mutations that give rise to the *TSG*^−/−^ genotype is given by *n*_2_ = 2. The mutation rate is denoted by *μ*. Finally, the system is characterized by the carrying capacity *K*, which describes the maximum number of cells that can exist in this system. We do not explicitly include the dynamics of the advantageous *TSG*^+/+^ cells in this model because the contact process simulations are stopped as soon as the first cell of this type is generated.

System (1-4) cannot be obtained by averaging the dynamics of the contact process, because of the presence of higher moments. The ODE’s however provide valuable insights into the system behavior.

In the subsequent analysis, we denote for convenience

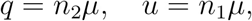

which are the total mutation rates of the passenger mutation production and TSG inactivation, respectively. There are 5 distinct steady states in this system, only one of which is of interest to us.

**Selection mutation equilibrium.** If *q* < 1 − *s*_3_, this model is characterized by a stable equilibrium that describes the coexistence of all cell populations. It is determined by a balance between the rate at which these cells are generated by mutation, and the rate at which they are lost as a result of their selective disadvantage (costs *s*_1_, *s*_2_, and *s*_3_). Hence, this level is referred to as the “selection-mutation balance”. The dominant populations are *x* and *z*, and they are given by

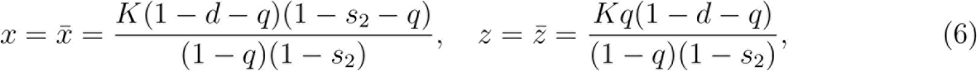

where terms of the order *u* and higher are omitted. The other two populations are of the order *u*:

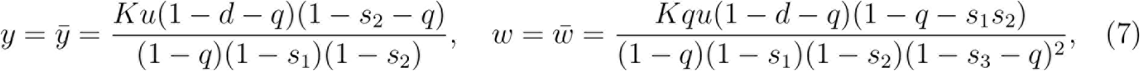

where terms of the order *u*^2^ and higher are omitted. At this equilibrium, the total population is given by

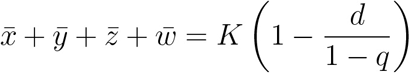

(again, only the zeroth order in *u* is kept).

**Double-hit mutant production.** The quantity of interest is the total rate of double-hit mutant (*TSG*^+/+^) production, given by

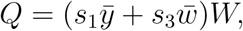

where the number of intermediate *TSG*^+/−^ cells is taken at the selection-mutation equilibrium. This quantity represent the total division rate of the two populations, *y* and *z*, capable of producing double-hit mutants; the factor *W* represents the effect of finite carrying capacity. We have

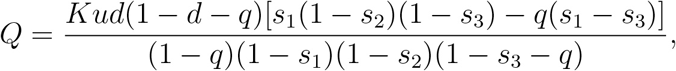

To determine whether passenger mutations facilitate the production of double-hit mutants, we consider the quantity *dQ/dq*. Positivity of this quantity indicates that a higher intensity of passenger mutant production will increase the rate of double-hit mutant production. We have

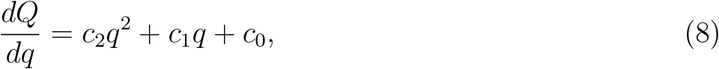

where coefficients *c*_0_, *c*_1_, *c*_2_ depend on *s*_1_, *s*_2_, *s*_3_, and *d*:

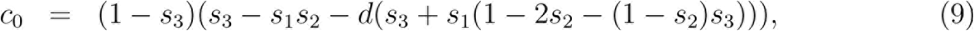

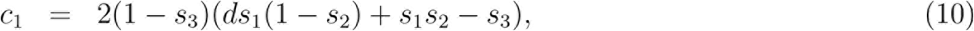

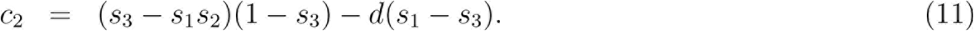

Depending on the signs of these coefficients, the behavior of the quadratic function (8) changes. All the cases are mapped out in figure 1 for two different parameter sets. The signs of the three coefficients are shown as regions in the *d* − *s*_3_ diagram, with *s*_1_ and **s*_2_* fixed to particular values. The three lines in figure 1 (a) and (b) correspond to *c*_0_ = 0, *c*_1_ = 0, and *c*_2_ = 0. It can be shown that all three lines intersect in one point, given by

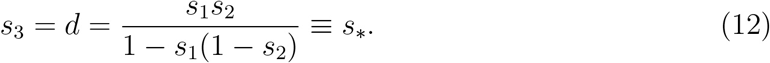

**Figure 1:**
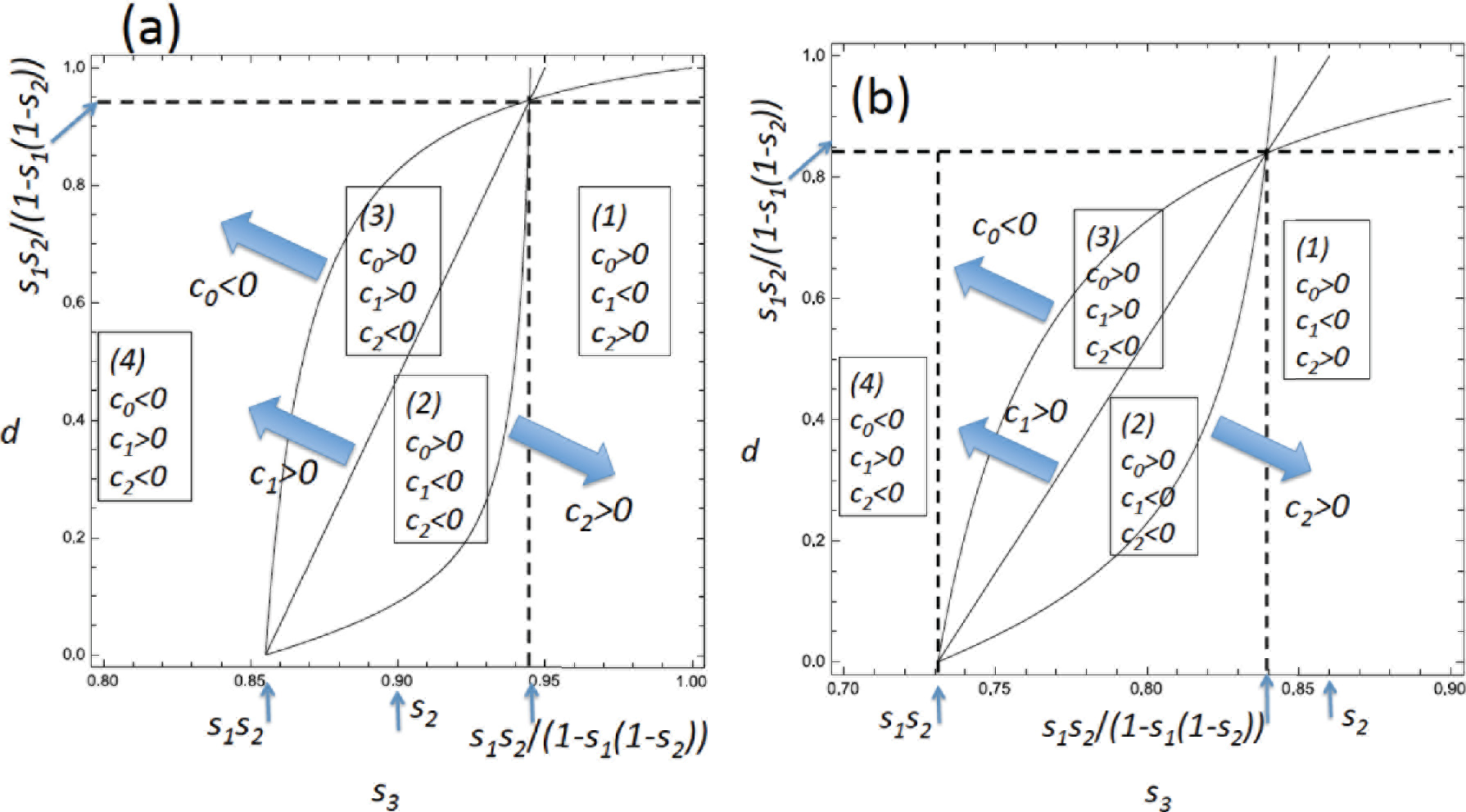
Phase diagrams showing the sign regions of parameters *c*_0_, *c*_1_, and *c*_2_, equations (8) and (9-11). The axes are *s*_3_ (horizontal) and *d* (vertical). The other parameters are fixed at (a) **s*_1_* = 0.95 and *s*_2_ = 0.9; (b) *s*_1_ = 0.86 and *s*_2_ = 0.85.

If *s*_1_ and *s*_2_ are close to 1, then this quantity is also close to 1, and it does not make biological sense to consider values of *d* larger than this threshold. Therefore, there are 4 biologically relevant regions in the *s*_3_ — *d* space, which are all marked in the figure.

Let us denote by *q*_1_ and *q*_2_ the smaller and the larger roots of the polynomial (8), respectively:

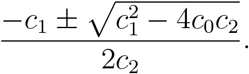

In the four different regions of figure 1, the quantity *dQ/dq* is positive in different intervals of the parameter q. These are listed below. Recall that *dQ/dq* > 0 is interpreted as “passenger mutations facilitate double-mutant production”.

(1) In this regime, see figure 2(1), *dQ/dq* > 0 for all values of *q* if

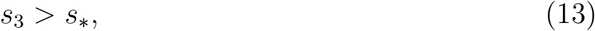

see definition (12). This regime is depicted by the upper parabola in figure 2(1), and by the horizontal line marked “(1) upper” in figure 3. Note that this regime is not always relevant. Biologically speaking, the fitness *s*_3_ must be bounded as

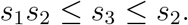

It may happen that *s*_*_ > *s*_2_, as in figure 1(a); in this case, this regime is biologically irrelevant. In general, we have

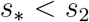

as long as

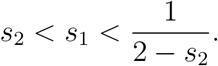

This inequality holds for the parameters of figure 1(b). The region of values allowed by this inequality for *s*_1_ shrinks as *s*_2_ grows. If inequality (13) is reversed (the lower parabola in figure 2(1)), that is,

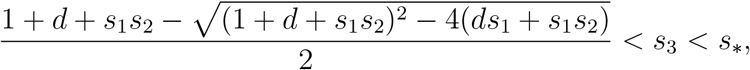

the condition for *dQ/dq* > 0 becomes

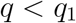

(note that *q*_2_ > 1 in this regime). This is illustrated by the horizontal line marked “(1) lower” in figure 3(a).

(2) and (3) These regimes are characterized by

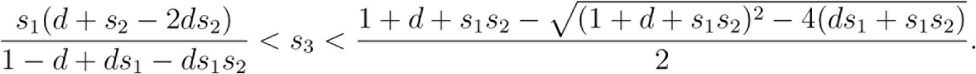

In these regimes, see figure 2(2) and (3), *dQ/dq* > 0 for all values of *q* if

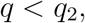

see also the horizontal line marked “(2) and (3)” in figure 3(a).

(4) Finally, we have the regime where

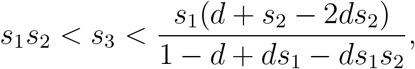

figure 2(4). In this case, we have

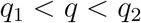

(see the upper parabola in figure 2(4) and the horizontal line marked “(4) upper” in figure 3(a)). In the biologically irrelevant case where *s*_3_ < *s*_1_*s*_2_, no values of *q* will correspond to *dQ/dq* > 0 (the lower parabola in figure 2(4) and the horizontal line marked “(4) lower” in figure 3(a)).

**Figure 2:**
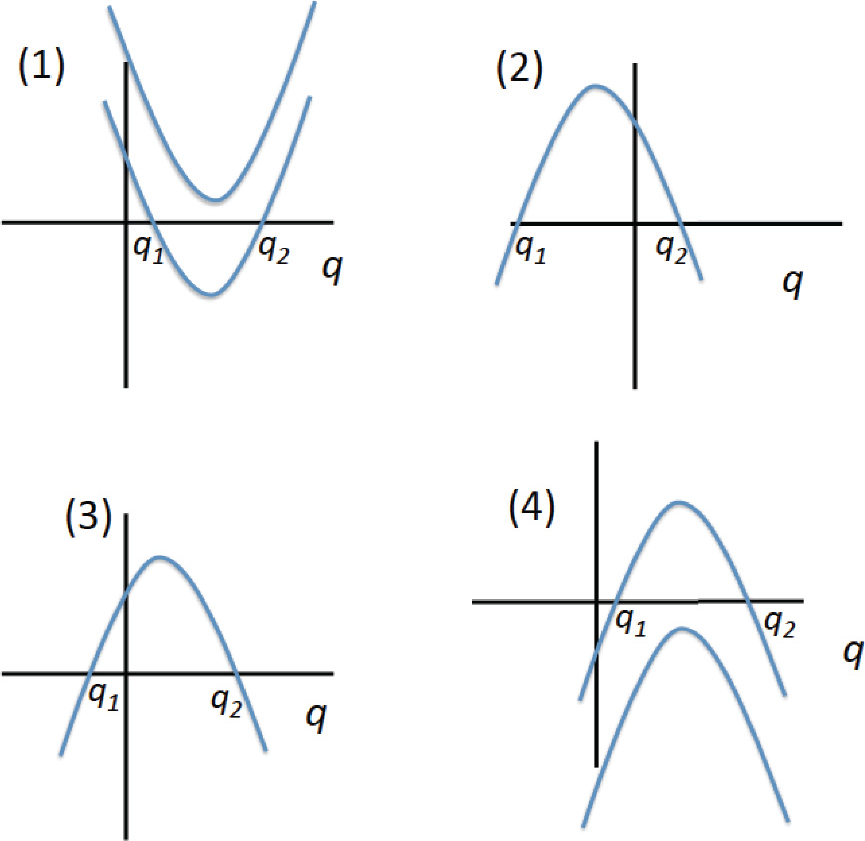
The four regimes identified in figure 1. For each regime, the parabola *dQ/dq* (equation (8)) is plotted schematically, to show the position of the relevant interval of *q*, depending on the signs of the coefficients *c*_0_, *c*_1_, *c*_2_.

**Figure 3:**
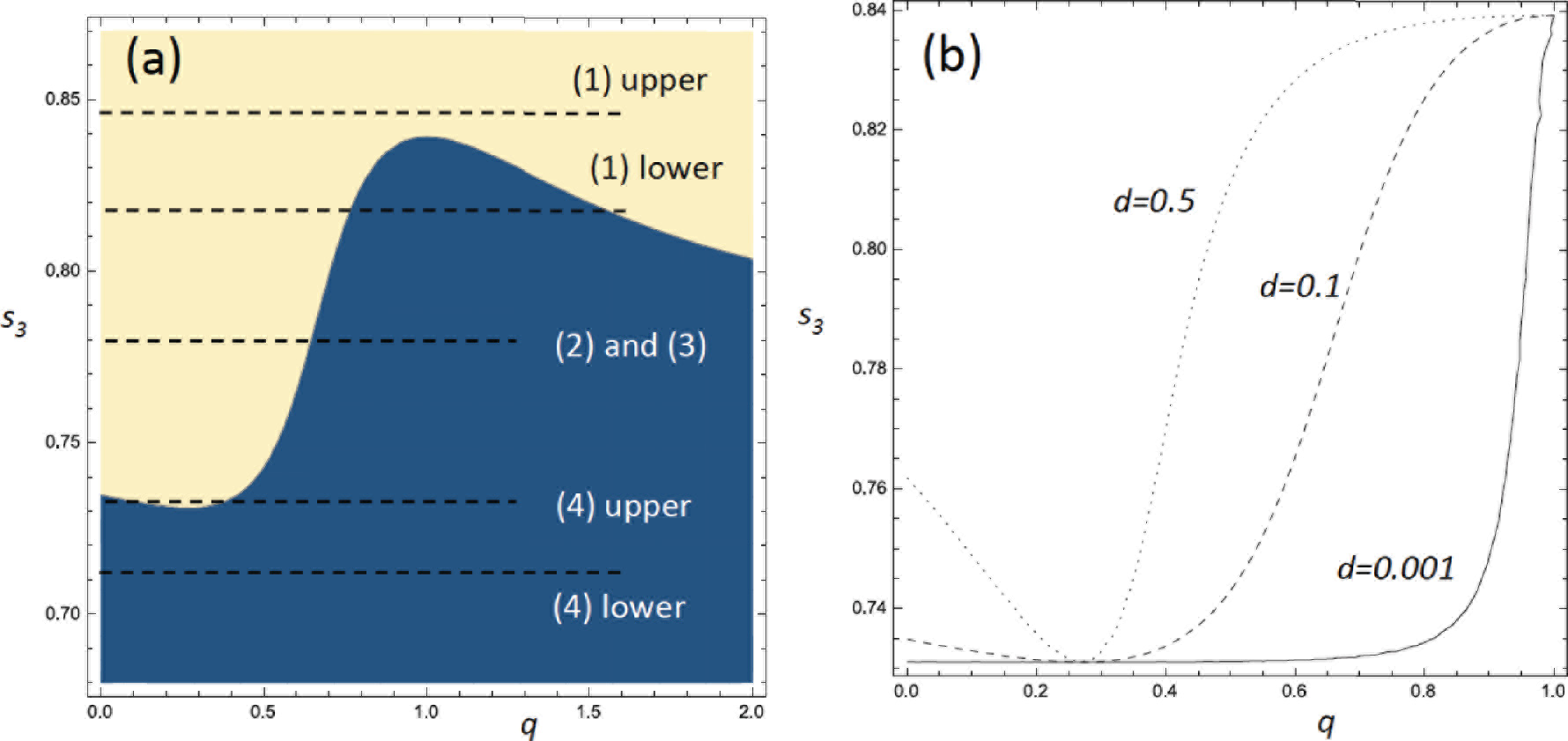
The *s*_3_ − q diagram showing the regions where *dQ/dq* is positive; the parameters correspond to figure 1(b). (a) For a fixed *d* = 0.1, the regions where *dQ/dq* > 0 and *dQ/dq* < 0 are shown in yellow and blue respectively. Dashed horizontal lines correspond to fixing a particular value of *s*_3_, in the four regimes of figure 1(b). (b) The contour lines *dQ/dq* = 0 for three different values of *d* (*dQ/dq* is positive above the lines).

The region where *dQ/dq* > 0 shrinks as *d* grows, as shown in figure 3(b), and the ODE model predicts that the effect of the passenger mutations weakens for larger *d*. On the other hand, the applicability of the ODE approach to explain the behavior of the stochastic system breaks down for large values of *d*, because as *d* grows relative to the division rates, the population size shrinks, and in the stochastic system, instead of the phenomenon of tunneling, one observes sequential fixation. As a consequence, we find that the passenger mutations play a stronger enhancing role in the creation of double-hit mutants.

### 1.2 Quasispecies equations

An alternative way to model the dynamics of the competing types in a fixed population is to use the quasi species type equations:

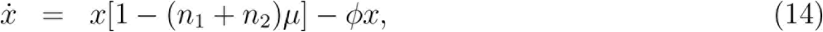

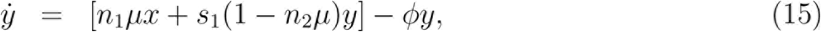

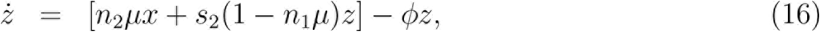

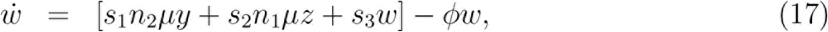

where

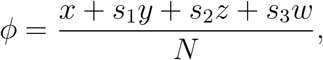

and *N* is the total, fixed population. The difference between this system and the previous one is the fact that in equations (14-17) the population always remains equal to *N*, that is, *x* + *y* + *z* + *w* = *N*, and the number of independent variables is effectively three, and the death rate (given by *ϕ*) is non-constant, as it adjusts to exactly balance the divisions. In equations (1-4), the death rate is fixed (to *d*), the population is non-constant, and all populations are independent variables. While equations (1-4) are analogous to the contact process, quasispecies equations can be thought as an ODE equivalent of the Moran process.

The mutation-selection equilibrium of this system is stable if *q* < 1 − *s*_3_, and is identical to that given by equations (6,7), where we set *d* = 0 and replace *K* → *N*. At this equilibrium we have

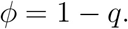

Note that although the equilibrium of the quasi species system is given by equations (6,7) with *d* = 0, it cannot be obtained from system (1-4) directly by setting *d* = 0 in the ODEs. Equations (1-4) with *d* = 0 do not describe the same dynamics as the quasi species system; one obvious difference is the absence of death in the former system, and the death rate given by 0 in the latter. Aside from the trivial equilibrium, equations (1–4) with *d* = 0 have a family of neutral equilibria that satisfy *x̅* + *y̅* + *z̅* + *w̅* = ***K***, with no further restrictions on the values of the individual populations.

Returning to the quasi species system, consider the function that defines the rate of double-hit mutant production. This is given by

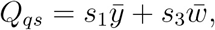

where the number of intermediate *TSG*^+/−^ cells is taken at the selection-mutation equilibrium. We have

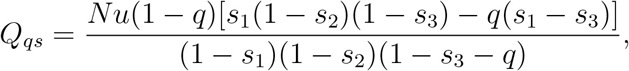

where terms of order *u*^2^ and higher are omitted. It is interesting to investigate how this quantity depends on *q*, the rate of passenger mutant production. Considering *dQ*_*qs*_/*dq*, we can see that this quantity can be positive only if *s*_3_ > *s*_1_*s*_2_, which, together with inequality (5) gives the following bounds for the quantity s_3_:

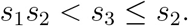

**Figure 4:**
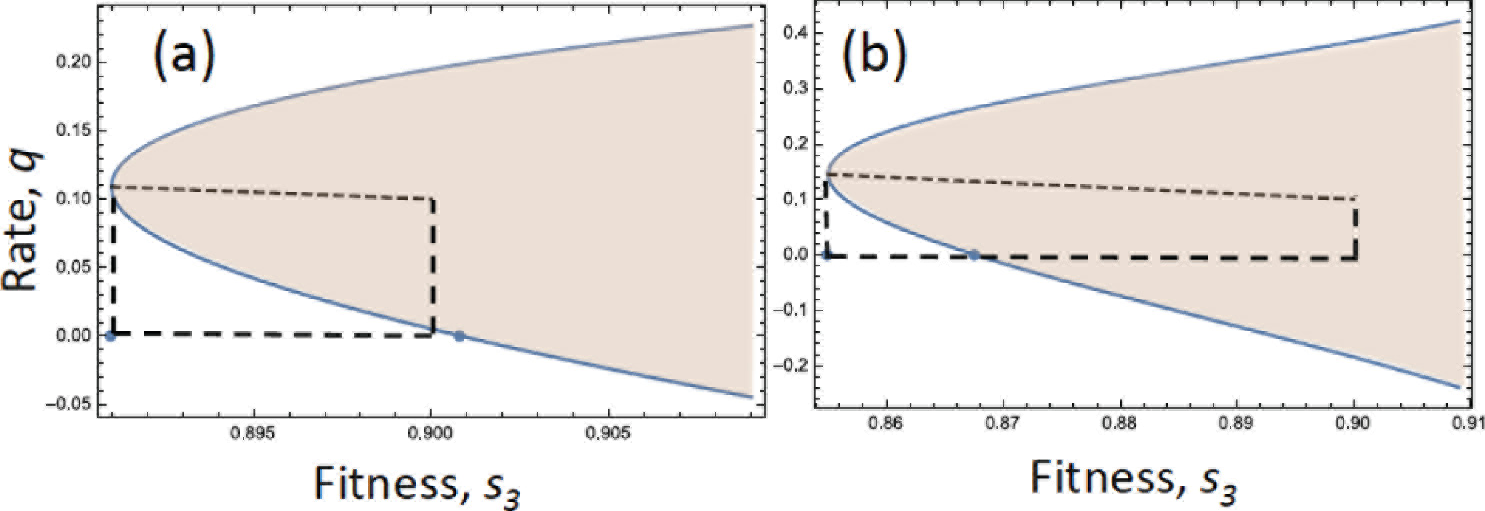
A parametric plot showing the dependence of the production term for the quasi species equations, *Q*_*qs*_, on the rate of passenger mutations, *q*. The horizontal axis is *s*_3_, with vertical dashed lines restricting this quantity to the range *s*_1_*s*_2_ ≤ *s*_3_ ≤ *s*_2_. The vertical axis is *q*, with dashed lines restricting this quantity to the range 0 ≤ *q* ≤ 1 − *s*_3_. The shaded regions correspond to the quantity *Q*_qs_ increasing with *q* (i.e. *dQ*_*qs*_/*dq* > 0); they correspond to condition (18). The parameters *s*_2_ = 0.9, and (a) *s*_1_ = 0.99; (b) *s*_1_ = 0.95.

Under this restriction, we have *dQ*_*qs*_/*dq* > 0 as long as

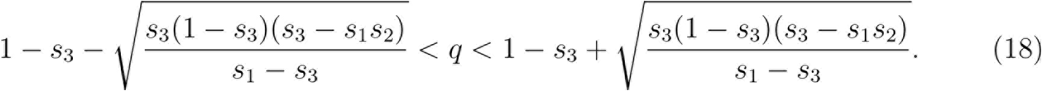

This is illustrated in figure 4.

## 2 The contact process: further details

### 2.1 Stochastic fitness of passenger mutants

The main analysis has been done by assigning passenger mutants an “average fitness”, defined by their division rate. In reality, however, different passenger mutations can bring about different costs to fitness. Therefore, we also constructed a model in which the fitness cost of the passenger mutants was drawn from a distribution with values between 0 (maximum cost) and 1 (no cost, the division rate is equal to that of the *x* population), and a mean of *s*_2_. Specifically, upon generation, each mutant cell was assigned a fitness value, given by *g*^1/*α*^, where *g* is a uniformly distributed random number in [0,1], and *α* = *s*_2_/(1 − *s*_2_). This corresponds to the beta distribution *B*(*α*, 1), which is a power function distribution on [0,1], which has mean *s*_2_. Similar methodology was used to generate passenger mutants with one copy of TSG inactivated (type *w*). As seen in Figure 5, the accelerating effect of passenger mutations remains robust in this model.

### 2.2 Spatially structured cell populations

We also investigated a model version which violated the assumption of mass-action. Thus, upon division, we assumed that the offspring cell could only be placed into the immediate vicinity of the source cell. To do so, we constructed a two-dimensional grid and tracked the location of each cell on the grid. The place for the offspring cell was randomly chosen from the eight nearest neighboring spots. If the chosen spot was empty, the daughter cell was placed there, otherwise, the division was aborted. Apart from that, the model remained the same. As seen in Figure 5, the accelerating effect of passenger mutations remains robust in the spatially structured model, although the extent of the acceleration is less pronounced in the spatial compared to the non-spatial model. The reason is that evolution of the double mutant in the absence of passenger mutations is accelerated in the spatial relative to the non-spatial model, while the rate of evolution is the spatial and non-spatial models is similar in the presence of passenger mutations. The effect of space on the rate of fitness valley crossing in this type of model is complex. Space can accelerate or slow down the rate of fitness valley crossing, depending on the exact parameter combinations. Therefore, in the spatial model, the effect of passenger mutations can in principle be either weaker than in the non-spatial model (as shown here), or it can be stronger. This has not been explored further here, because it is beyond the scope of the current investigation.

## 3 Moran process

As explained in the main text, the assumptions underlying the contact process were also formulated as a Moran process, which assumes a constant population. We analyzed the model assuming that passenger mutants are characterized by an average fitness cost, *s*_2_, and that cells can mix perfectly (mass action). The Moran process was used for the following reasons: (i) This model is more amenable to mathematical analysis, which is presented below. (ii) The constant population Moran process is more suited to describing the evolutionary dynamics in healthy tissue, which is tightly regulated by homeostatic mechanisms. While the occurrence of passenger mutations is mostly discussed in the context of a cancer, the literature documents an array of seemingly non-functional mutations in the healthy tissue of people, especially with advancing age [6, 11].

**Figure 5:**
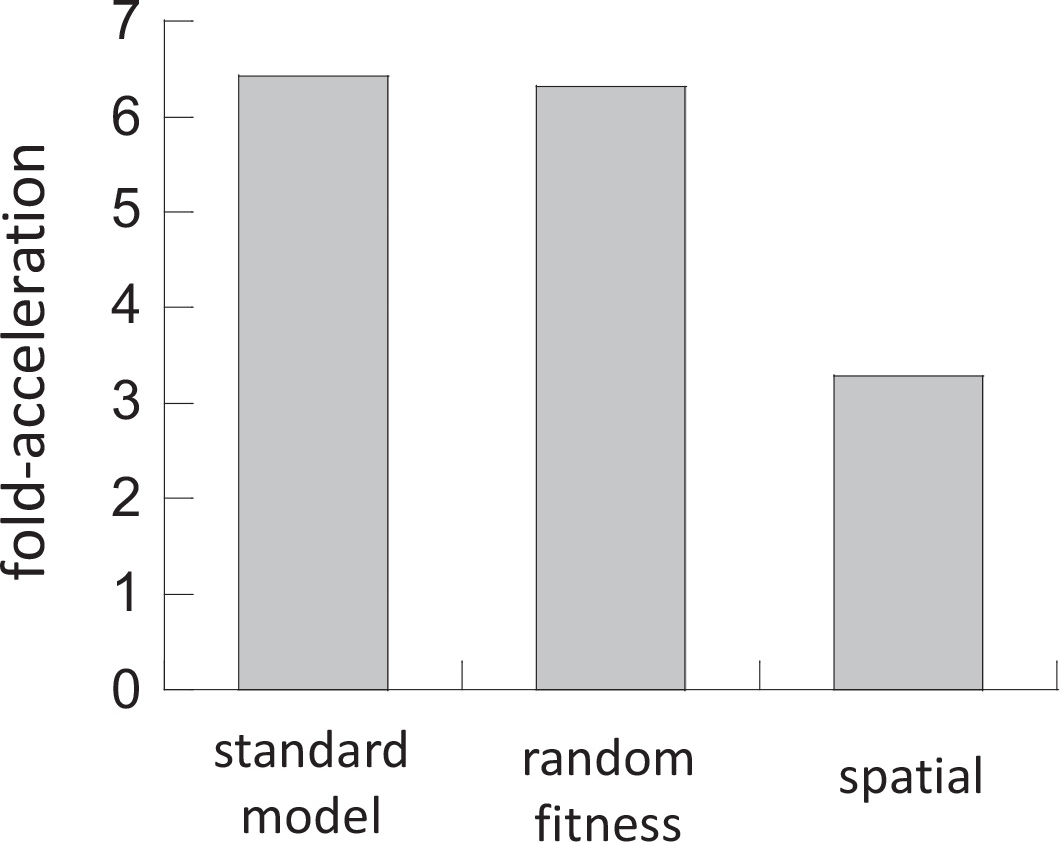
Effect of alternative model assumptions. The fold-acceleration of evolution due to passenger mutants is plotted for different models. The computer simulations were run repeatedly, and the fraction of realizations in which the double mutant was created before a time threshold was determined. The fold acceleration is the fraction of runs where the double mutant was generated in the presence of passenger mutations divided by the fraction in the absence of passenger mutations. This is done for one specific value of passenger mutant fitness, *s*_2_/*s*_3_. The standard model corresponds to the model explored in the main text. The random fitness models draws passenger mutant fitness from a distribution as explained in the text of the supplementary materials. The spatial model assumes that cell offspring can only be placed into the eight nearest neighboring spots, as explained in the text of the supplementary materials. All three fold increases are statistically significant by the Z-test (*p* < 0.05). Parameters were chosen as follows. *L*_x_ = 0.15, *D* = 0.01, *μ* = 10^−6^, *s*_1_ = 0.99, *s*_2_ = 0.99, *s*_3_ = 0.99, *n* = 5 × 10^−2^/*μ*, *T*_thr_ = 120,000, *N* = 2500.

### 3.1 Numerical simulations

The accelerating effect of passenger mutations is the same as that observed in the contact process, and the parameter dependencies remain qualitatively the same as in the contact process. Figure 6 repeats the analysis and shows the same plots as done for Figure 2 in the main text.

A mathematical analysis of the Moran process is presented next.

### 3.2 No passenger mutations

We envisage a Moran process where a constant population of *N* cells is in a steady turnover. At each elementary update, each cell has the same probability to die (given by 1/*N*), and then this cell is replaced by an offspring of another cell. Cells are chosen for reproduction with a probability proportional to their fitness value. One unit of time corresponds to *N* elementary updates. We will investigate a general process where three types of mutations are possible. The first mutation (which corresponds to the inactivation of the first copy of a TSG) happens with probability *u*_1_ per cell division. The second copy of a TSG is inactivated with probability *u*_2_. Finally, passenger mutations are acquired with probability *nu*.

Denote the number of *TSG*^+/−^ mutants of fitness *s*_1_ by *i*, *i* ∈ {0,1, 2,…}, and by *φ*_i_(*t*) the probability to have *i* mutants at time *t*. State *E* corresponds to the creation of the first double-hit mutant, and this is an absorbing state. We have:

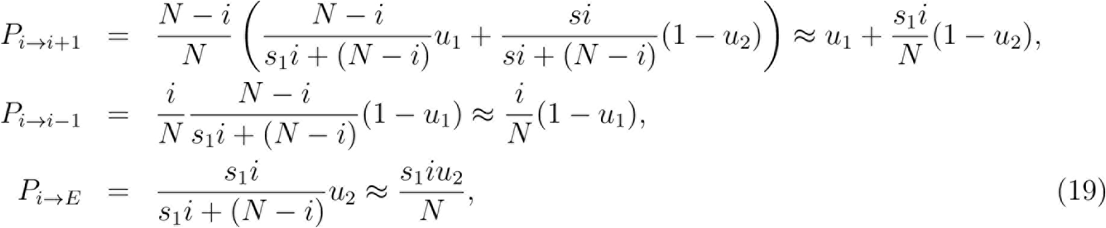

where we neglected *i* compared to *N*. We will rescale time by having *N* elementary events during a unit time. The function *φ*_i_(*t*) satisfies the following Kolmogorov forward equation,

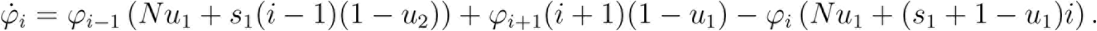

Introducing the probability generating function,

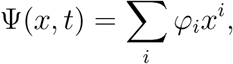

we can derive the following 1st order PDE:

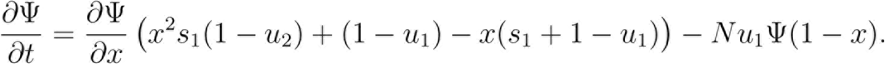

Suppose *x*(*t*) is the solution of the initial value problem,

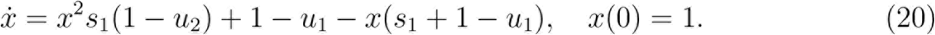

**Figure S6:**
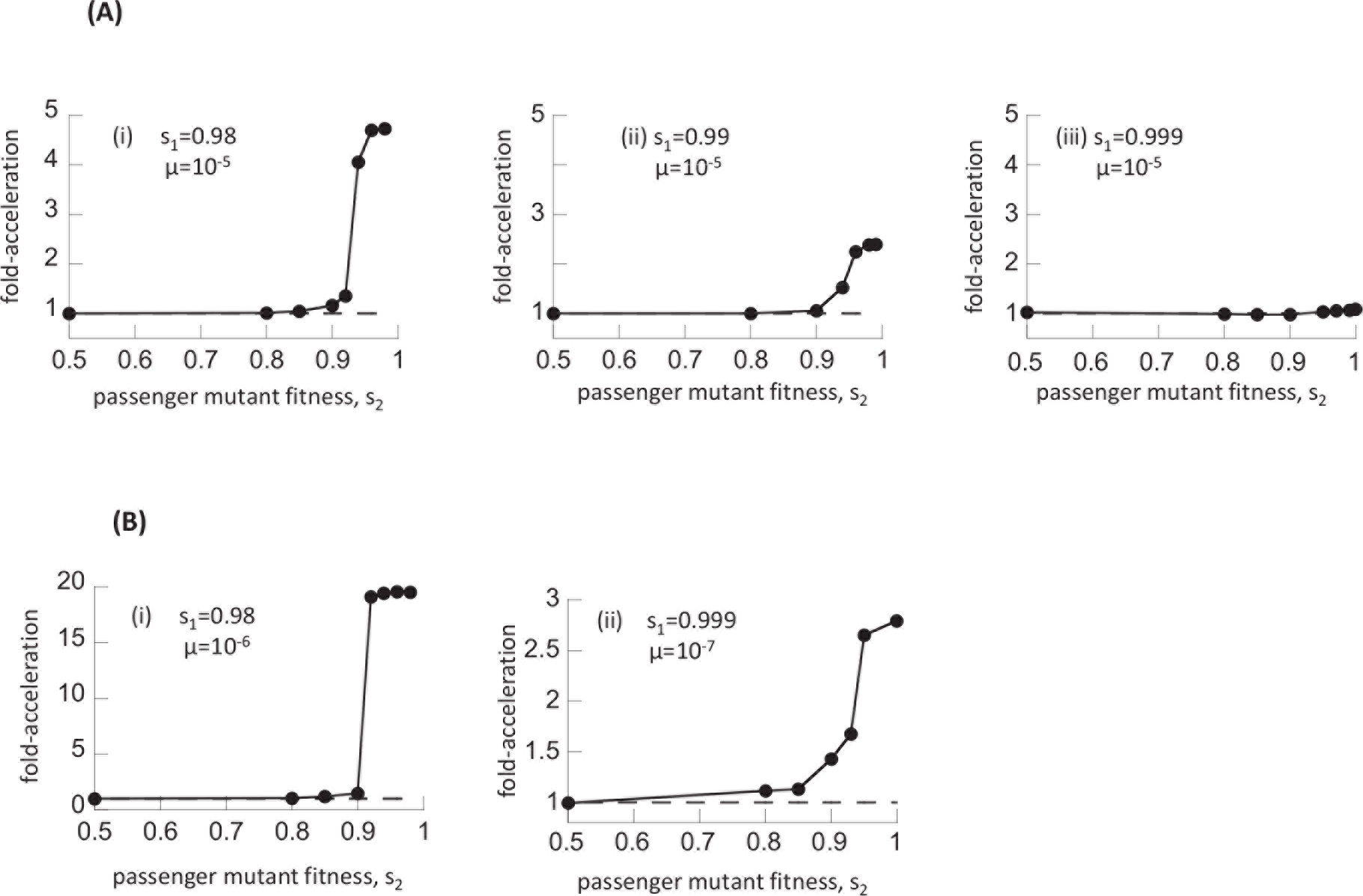
Accelerated evolution through passenger mutations in the Moran process model. The computer simulation was run repeatedly, and the fraction of realizations in which the double mutant was created before a time threshold T*thr* was determined. Each graph plots the “fold acceleration”, which is the fraction of runs where the double mutant was generated in the presence of passenger mutations with fitness *s*_2_, divided by the same measure in the absence of passenger mutations. (A) Parameter regime where the double mutant evolved through a tunneling pathway. Panels (i-iii) show that a reduced cost of the intermediate TSG^+/−^ mutant (higher value of *s*_1_) leads to a reduced effect of passenger mutations on the rate of evolution. (B) Parameter regime where the double mutant evolves by sequential fixation, in which passenger mutants have a stronger accelerating effect on the rate of evolution. Parameters were chosen as follows. We assumed *s*_3_ = *s*_2_, although results do not depend on this constraint, see above; *n* = 5 × 10^−2^/*μ*. The total population size was *N* = 2500. The time threshold are given as follows. (A) (i) *T*_*thr*_ = 5, 000; (ii) *T*_*thr*_ = 500; (iii) *T*_*thr*_ = 200. (B) (i) *T*_*thr*_ = 50, 000; (ii) *T*_*thr*_ = 50, 000. Differences between outcomes with and without passenger mutations were statistically significant (*p* < 0.05) for A(i) *s*_2_ ≥ 0.85, A(ii) *s*_2_ ≥ 0.9, A(iii) *s*_2_ ≥ 0.95, B(i) *s*_2_ ≥ 0.85, B(ii) *s*_2_ ≥ 0.8.

The probability to create a double-hit mutant by time *T* is given by

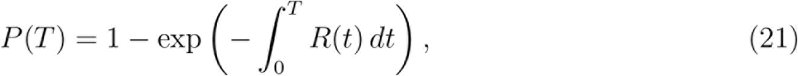

where the tunneling rate is given by

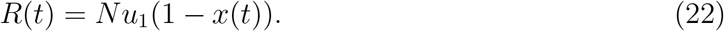

### 3.3 Including the possibility of passenger mutations

Consider a Moran process. We assume that *n* ≫ 1, such that the rate of passenger mutations is much larger than the TSG mutation rate. Let us ignore the TSG mutations, and find the population equilibrium. This can be done by considering the quasi-species equations:

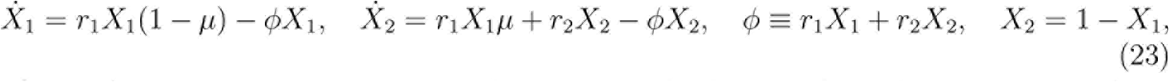

where *μ* is a one-way mutation rate and *r*_1_ > *r*_2_ are the fitness values of the two populations, *X*_1_ and *X*_2_, respectively. We have

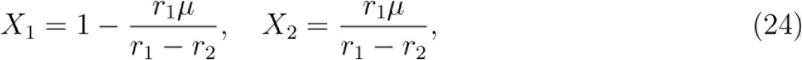

as long as 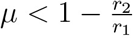; otherwise we have *X*_1_ = 0 and *X*_2_ = 1.

Populations *x* and *w* have the fitness values of 1 and *s*_2_ respectively, and the mutation rate is given by *nu*. Therefore, we can assume that at a quasi-steady state, we have a population of wild type cells of size *x* = *N* − *j*_0_, and the population of cells with passenger mutations, *w* = *j*_0_, given by

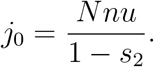

The mean fitness of this “resident” population is given by

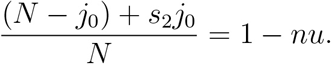

We further assume that the following processes happen in the system:

- population *w* acquires passenger mutations, population *z*, at rate 2*u*.
- population *x* creates mutants *y* at rate 2*u*,
- population *z* creates double-hit mutants at rate *u*_2_,
- population *y* creates double-hit mutants at rate (1 − *nu*)*u*_2_.

By following the same method as before, we denote the number of mutants of fitness *s*_1_ by *i*, and the number of mutants of fitness *s*_3_ by *j*, with *i*, *j* ∈ {0,1, 2,…}. The function *φ*_*i,j*_ (*t*) is the probability to have *i* and *j* mutants of each type at time *t*. As before, state *E* corresponds to the creation of the first double-hit mutant, and this is an absorbing state. We have:

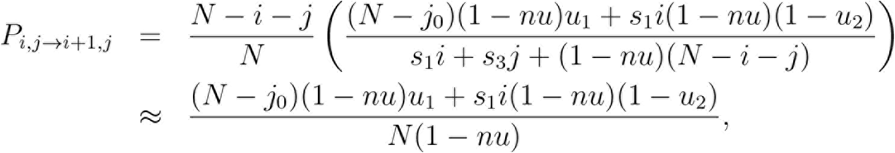

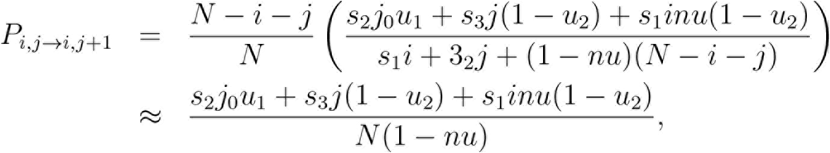

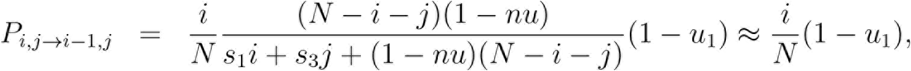

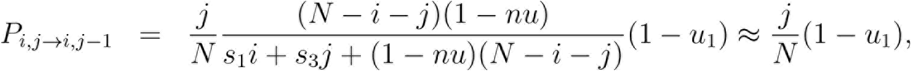

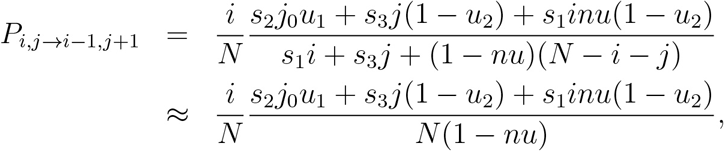

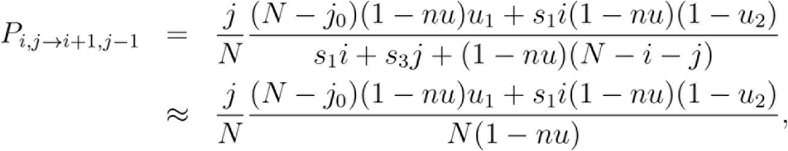

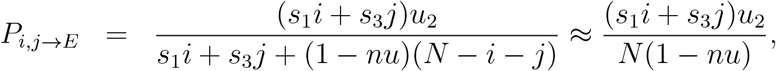

where we neglected *i* and *j* compared to *N*. We will rescale time by having *N* elementary events during a unit time. The function *φ*_*i,j*_(*t*) satisfies the following Kolmogorov forward equation,

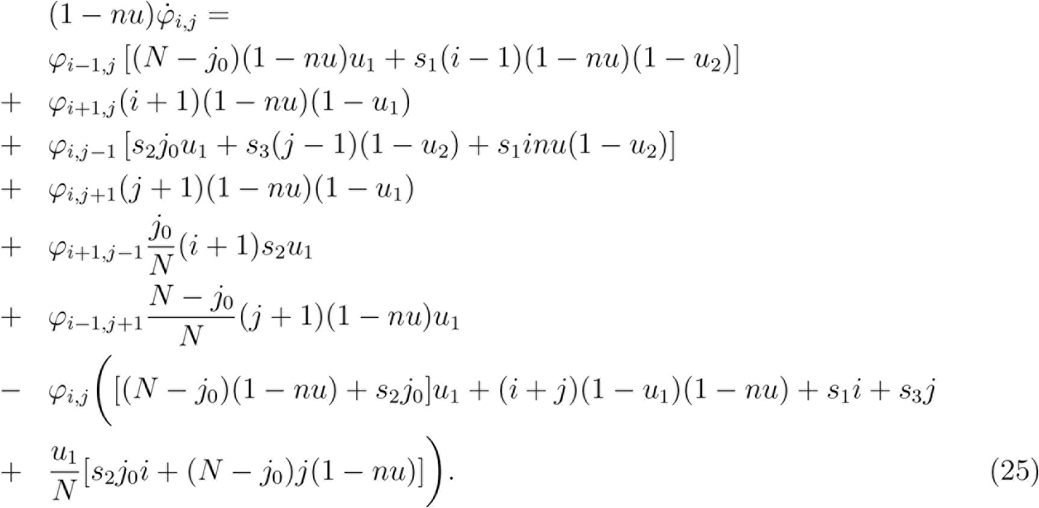

In the above equation, higher order terms (nonlinear in *i/N*, *j/N*) are omitted. Introducing the probability generating function,

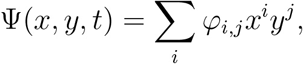

we can derive the following 1st order PDE:

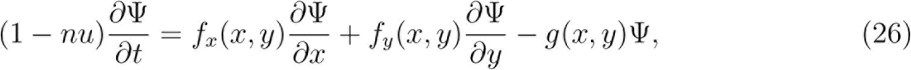

where

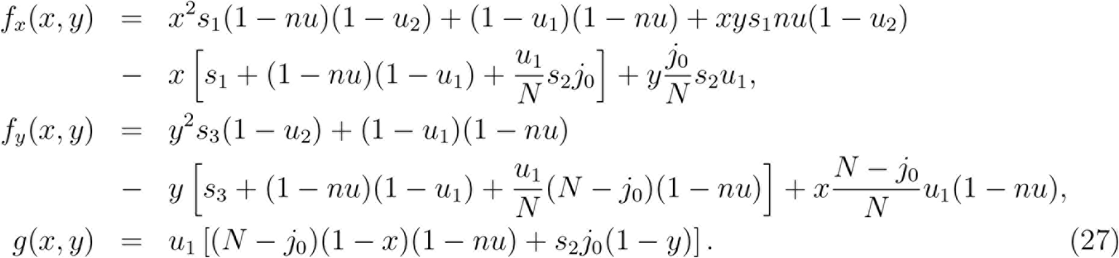

Suppose that *x*(*t*) and *y*(*t*) are the solutions of the initial value problem,

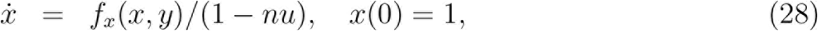

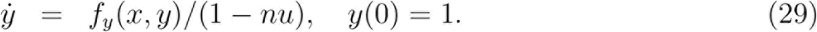

Then the probability to create a double-hit mutant by time *T* is calculated as

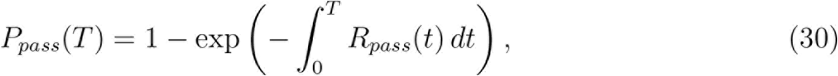

where the tunneling rate, *R*(*t*), is given by

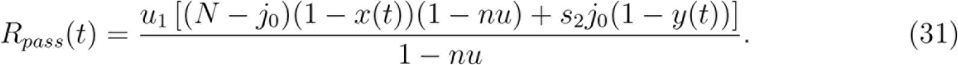

We can see that if *n* = 0 (no passenger mutations), we have *j*_0_ = 0 and *R*(*t*) = *R*_*pass*_(*t*), that is, equations (22) and (31) coincide, as they should.

### 3.4 Comparisons

Figure 7 summarizes the theoretical results and compares them with the results of stochastic simulations. For simplicity we assumed that the fitness of *TSP*^+/−^ cells is *s*_2_, irrespective of their passenger mutation status; similar results will hold for a range of assumptions on fitness *s*_3_. The fitness of *TSG*^+/−^ cells in the absence of passenger mutants, *s*_1_, was fixed, and several values of parameter *s*_2_ = *s*_3_ were tested. For each value of *s*_2_, a large number of simulations were run. Each simulation was stopped when either a *TSG*^−/−^ mutant was created, or a stopping time *T* was reached. For each simulation, we recorded whether a *TSG*^−/−^ mutant was created, and then the probability of *TSG*^−/−^ creation by time *T* was calculated. This is shown by blue dots in figure 7. Thes numerical results were compared with quantity *P*_*pass*_, equation (30), which is plotted as a red curve in figure 7. The horizontal lines in this figure correspond to the numerically calculated (blue) and analytically obtained (red, formula (21)) values *P*(*T*), the probability to generate a *TSG*^−/−^ mutant in the absence of passenger mutations.

**Figure 7:**
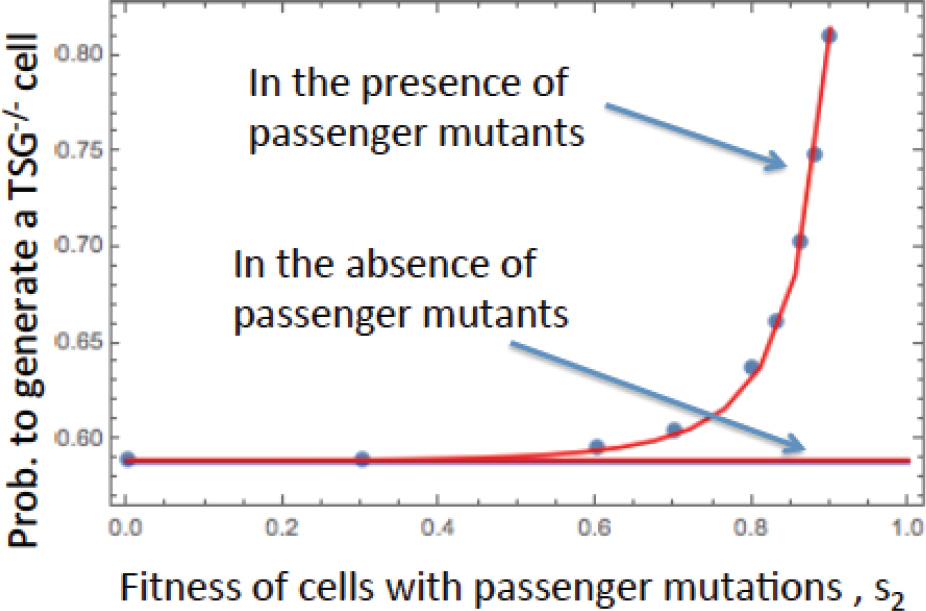
The probability to create a TSG^−/−^ mutant by time *T*. Numerical simulations are depicted in blue, and analytical results (formulas (21) and (30)) in red. The parameters are *s*_1_ = 0.9, *u* = 10^−4^, *u*_1_ = 2*u*, *u*_2_ = *u*, *n* = 500, *T* = 2000. The number of simulation was more than 500, 000 for each value of *S*_2_.

First of all, we observe that numerical and analytical results are in a very good agreement. Secondly, we can see that the probability of *TSG*^−/−^ mutant creation is higher in the presence of passenger mutation, as long as the fitness of the corresponding cells is not too low (but lower than the base fitness of single hit mutants, *s*_1_).

## 4 Connections between the role of 3D perturbations in turbulence/ patterns and passenger mutations in 2-hit mutant generation

The idea that initially disadvantageous or passenger mutations might allow cells make a more rapid transition to carcinogenesis by providing more effective pathways for normal cells to reach that state stems from analogies with the onset of turbulence in fluids and the nucleation of dislocation defects in natural patterns.

No one area of science stands alone. Analogies can provide useful insights. Behavior seen in one context can be seen in another where it initially it might not have been expected. The common denominator is often mathematics. For example, freak waves, about which ocean navigators have many terrifying stories and experiences, also occur, surprisingly perhaps, in optical fibers. Their common origin is modulated wave-trains whose behaviors are captured in water waves, in optics, in plasmas by non-integrable corrections (in the water wave context, the full Euler equations) to the canonical and universal nonlinear Schroedinger equation for wave envelopes. In the right circumstances, the slowly modulated wave-train can evolve into a signal in which one sees a burst of very large amplitude waves over very few wavelengths. The phenomenon is robust and universal.

The analogies which motivated the present studies had their origins in the onset of turbulence and in pattern formation. The study of the onset of turbulence in shear flows goes back to the classical experiments of Osborne Reynolds in 1883 on pipe flow in which water flows along a pipe [14, 15]. He found that typically the flow will be in the turbulent state at values of the Reynolds’ number, *UD*/*ν*, of about 2000 and laminar below that. *U* is the flow velocity say on the center line, *D* the pipe diameter and *ν* the kinematic viscosity. If he was very careful to avoid creating disturbances at the inflow point (he could use a funnel to minimize these), then he, and others later, showed that one could retain the laminar character of the flow up to Reynolds numbers of over 10,000. The conclusion is that turbulence onset in pipes is a nonlinear and three dimensional phenomenon and requires finite amplitude perturbations to trigger the turbulent state.

That conclusion is supported by the negative results emanating from linear stability theory which, for shear flows such as pipe flow, plane Poiseuille flow (pressure driven flow in a channel), Couette flow (flow between oppositely moving plates), boundary layer flow (flow over a flat plate) does not predict what is observed. For both pipe and Couette flow, the flows are linearly stable for all Reynolds numbers. For Poiseuille flow, linear instability sets in at *R* ~ 6000. In all cases, the observed onset of turbulence typically occurs at values of R much less than what linear stability predicts. Moreover, the observed turbulent fields have a richer character than do the original shear flows. The latter usually have the form (*U*(*y*), 0, 0), namely a velocity in the *x* or streamwise direction depending on one of the spanwise directions (the radial coordinate in pipes, the vertical coordinate in Couette, Poiseuille and boundary layer flows). On the other hand, turbulent flow depends on all three coordinates, *x*,*y* and *z*. Linear stability theory also gave rise to another misleading result. A theorem, due to Squire [16], says that of all the disturbances, the two dimensional ones are the least damped and, in the case of Poiseuille flow for *R* > 6000, would be the most amplified. So surely, if they were the least damped, they would also be the most important. Clearly that is not what is observed. Turbulence is strongly three dimensional. So linear stability theory does not necessarily identify the right set of shapes from which a turbulent field grows.

Such a conclusion was very evident in the early experiments of [7] (and earlier papers and colleagues) who demonstrated that turbulence is clearly a three dimensional phenomenon. Squire’s theorem pointed in the wrong direction. A theoretical breakthrough came with the work of [1] who discovered that three dimensional spanwise oscillations could give rise to shapes which grew algebraically in time. Such behavior could lead to a redistribution of streamwise velocity via what Marten Landahl called “up-lift”. Start with a two dimensional *x*,*y* perturbation with its vorticity (the swirl of the fluid) in the *z* direction. Think that the fluid rotates around the vortex line in a clockwise direction, down in front, up in back. Now perturb that vortex line, think cos(*bz*), so that near the front of this loop at *z* = 0, there is a flow which lifts fluid parcels up. The slower moving fluid which is carried away from the boundary into the faster moving flow outside can have a significant effect on the velocity profile *U*(*y*) and even can produce points of inflexion where *U*″(*y*) = 0. In boundary layers, such behavior can lead to very fast instabilities and turbulent bursts. Variations on this theme, namely the presence of algebraic in time growth of disturbances which are very weakly exponentially damped (have a t dependence *F*(*R*)*t* * exp(−*at*), *F* depends on the Reynolds number *R*, *a* is very small) which can lead to amplitudes at which linear and nonlinear terms in the flow equations are of equal importance, have been discussed in many papers in the past forty years. We list some notable ones: [1, 4, 9, 2, 13, 3, 17, 18, 19, 20, 5]. Trefethen et al identified the source of *t* growth multiplied by weak exponential decay behavior as generic if the linear stability operator *L* is non-normal. That is the case for shear flows. In short, the message is that comparing the relative and very slow exponential damping rates of different structures and then choosing the least damped or most amplified state as the one which controls the subsequent dynamics is not always correct.

In an altogether different context, similar behavior occurs when a pattern adjusts from one wavelength to another. Let *w* = *A* cos(*kx* + *f*) sin *z*, *A* exp(*if*) is called the complex amplitude, be the vertical velocity field of a set of roll-like cells whose axes are along the *y* direction. If the wavenumber *k* is too much larger than some preferred value *k*_0_, then an instability, called the Eckhaus instability, occurs, the net result of which is that the pattern will remove roll pairs until the new wavenumber is inside the Eckhaus stability boundary. The fastest growing linear instability mode has the envelope *A* remain a function of *x* and *t* only. Along the axis, that is *y* dependent, perturbations have a lower growth rate. Now in order to remove a roll pair with an envelope *A* only depending on *x*, the amplitude *A* has to enter its nonlinear regime and become zero along the full length of one pair of rolls. And, if one is very careful to control and suppress *y* dependent perturbations, that is indeed what happens [10]. But in most cases, and especially near or just inside the Eckhaus stability boundary, *y* dependent perturbations, which initially grow more slowly or decay slightly faster, overtake the *y* independent perturbation and dominate the outcome. Instead of having to remove a roll pair by making the real amplitude zero along the length of the roll, the *y* dependent perturbation produces a pair of dislocations at which positions the complex field *A* exp(*if*) is zero. The dislocations repel each other and travel to the boundaries at the end of the roll axes where they are absorbed, thereby removing an excess roll pair. It is easy to see that this scenario is a much more efficient means of wavenumber adjustment. The three dimensional disturbance, although less favorable according to linear stability analysis, is the one which eventually prevails. In short, allowing for three dimensional perturbations, the flow is deformed so that it can find a more efficient pathway to remove the roll pair.

The lesson we draw from these examples is this. Just because a structure is deemed disadvantageous with respect to others based on linear stability analysis does not mean that when the full dynamics is in play, it does not play a significant role. Whereas the structure may initially appear to be less fit, it may provide for a richer set of pathways in order that a system can achieve some desired goal more readily. Motivated by these ideas, we ask in this paper whether similar ideas might apply in enabling cells to use slightly disadvantageous mutations, here called passenger mutations, in order to provide a higher probability in reaching some genuinely malignant mutation by providing the deformed cell with a richer set of pathways to reach that goal.

